# Distinct antischistosomal profiles of antimalarials and TRP agonists

**DOI:** 10.64898/2026.10.29.761559

**Authors:** Nicolas J. Wheeler, Mostafa Zamanian, John D. Chan

## Abstract

Schistosomiasis control relies on praziquantel mass drug administration, but reports of field treatment failure and laboratory-selected resistance highlight the need for alternatives. Praziquantel activates a schistosome transient receptor potential (TRP) channel, TRPM_PZQ_. Alternative leads or chemotherapies include meclonazepam, which activates a distinct schistosome TRP channel, and the antimalarials artemether and mefloquine, whose flatworm targets are unknown. To compare molecular responses and assess cross-resistance, we conducted comparative RNA-seq and efficacy studies in infected mice. We profiled adult *Schistosoma mansoni* recovered 48 hours after artemether or mefloquine treatment and compared responses with existing datasets for TRP agonists praziquantel and meclonazepam. Transcriptional profiles clustered by drug class but showed largely distinct changes, with only 33 upregulated and 63 downregulated transcripts shared across treatments. Plexin domain-containing genes were strongly induced by both TRP agonists and, to a lesser extent, by antimalarials. Artemether downregulated gene families expressed in intestinal cells (saposins, cathepsins, and Bax inhibitor-like proteins), consistent with potential heme-dependent activity in the schistosome gut. Genes involved in glutamine metabolism, iron homeostasis and responses to oxidative damage were upregulated across multiple treatments. We evaluated all four treatments in mice infected with praziquantel-susceptible or resistant parasites. Praziquantel (200 mg/kg) reduced resistant male worm burdens by only 39.3%, versus 99.0% in susceptible parasites. Meclonazepam (83.0% versus 91.6% in susceptible parasites), mefloquine (99.1% versus 98.8%), and artemether (98.2% versus 99.4%) retained activity. The distinct transcriptional responses and retained activity against praziquantel-resistant parasites support further investigation of these compound classes as praziquantel alternatives or potential combination partners.

## Introduction

Blood-dwelling *Schistosoma* flatworms cause schistosomiasis, which is largely controlled by praziquantel (PZQ) mass drug administration. These programs have reduced the disease burden (1), but widespread use of a single drug risks selecting for resistance. PZQ-resistant schistosomes can be readily selected in the laboratory (2–5), and field studies over several decades have reported communities with low cure rates (6–8) and infections that remained uncured after three rounds of high-dose PZQ (9). Apparent resistance has also been reported in veterinary flatworm parasites, where PZQ use is widespread (10–12). Given these precedents, PZQ resistance emerging in the field with continued mass drug administration selection is a plausible concern. Additional antischistosomal drugs could provide alternative therapies when PZQ treatment fails or serve as partners in combination regimens intended to slow the emergence of resistance. This strategy is well established for malaria control (13) and has been proposed for schistosomiasis (14). In these scenarios, a key requirement is the absence of cross-resistance. Candidates intended to complement or replace PZQ should remain active when PZQ efficacy is reduced, and this is most likely when partner drugs act through distinct targets or pathways.

Promising alternatives to PZQ include certain benzodiazepines (15–17) and antimalarial drugs, such as mefloquine (18) and artemisinin-derivatives (19–21). PZQ is an agonist of the schistosome transient receptor potential (TRP) Ca^2+^ channel (Smp_246790) referred to as TRP_PZQ_ (5, 22). Benzodiazepines are thought to act on a different TRP channel (Smp_333650), referred to as TRP_MCLZ_ (23, 24). Although initially developed as antianxiety or antiseizure medications, some benzodiazepines were also found to kill schistosomes. The best studied lead compound, meclonazepam (MCLZ), has demonstrated efficacy in rodent models (15) and a small clinical trial (16). The schistosome target(s) of antimalarial drugs are less well understood. However, both intramammalian stages of schistosomes and blood-stage *Plasmodium* parasites digest host hemoglobin. Therefore, these drugs may have shared mechanisms in the two parasites centering around hemoglobin processing, heme detoxification, and redox stress.

The rationale for use of a drug as a PZQ alternative or combination therapy is strengthened if it has a distinct mechanism of action. In vitro phenotypes alone provide limited mechanistic insight and are often described using broad categories such as paralysis or reduced motility. Moreover, host immune responses contribute to the in vivo efficacy of many antiparasitic drugs. Here, we evaluated the effects of in vivo drug treatment on patent *Schistosoma mansoni* infections in a murine model of schistosomiasis, using RNA-Seq as a detailed, comparative readout of parasite response to drug exposure. Previously, we profiled the transcriptomic response of these parasites to the TRP agonists PZQ and MCLZ (25, 26). We reanalyzed these datasets using the updated genome assembly and annotations, and compared them with new transcriptomic datasets generated after treatment with two antimalarial compounds -mefloquine (MEF) and artemether (ART). These comparisons revealed unique profiles associated with each drug, implicating different cell or tissue types, together with changes shared across treatments that may reflect a common stress response or immune attack following drug-induced damage.

Finally, we have investigated the in vivo antiparasitic activity of these different drug classes against populations of *S. mansoni* that are susceptible or resistant to PZQ (5). We found no statistically significant evidence of cross-resistance with PZQ -MCLZ, ART, and MEF each retained activity against both PZQ-susceptible and PZQ-resistant parasites. Together, these data show that TRP-activating and antimalarial drugs produce distinct parasite responses and that reduced PZQ susceptibility does not necessarily predict reduced activity of these other drug classes. These findings support the study of mechanistically distinct compounds as PZQ-alternatives or potential partners for combination therapy.

## Results

### Comparative transcriptional responses to antischistosomal drug treatments

Antischistosomal drugs elicit different phenotypes and act on different timescales, consistent with different mechanisms of action. For example, TRP channel agonists (PZQ and MCLZ) cause contractile paralysis within seconds in vitro and parasite death in vivo within hours (25–28). In contrast, antimalarial drugs (ART and MEF) kill schistosomes, but with less obvious in vitro phenotypes and slower in vivo activity over at least two to three days (18, 19) (**Figure 1A**). To characterize the transcriptomic responses to these drugs, we compared parasite transcriptomes at time points associated with the hepatic shift of worms from the mesenteric vasculature to the liver, a characteristic response to anthelmintic treatment in the murine model of schistosomiasis (29).

**Figure 1.**
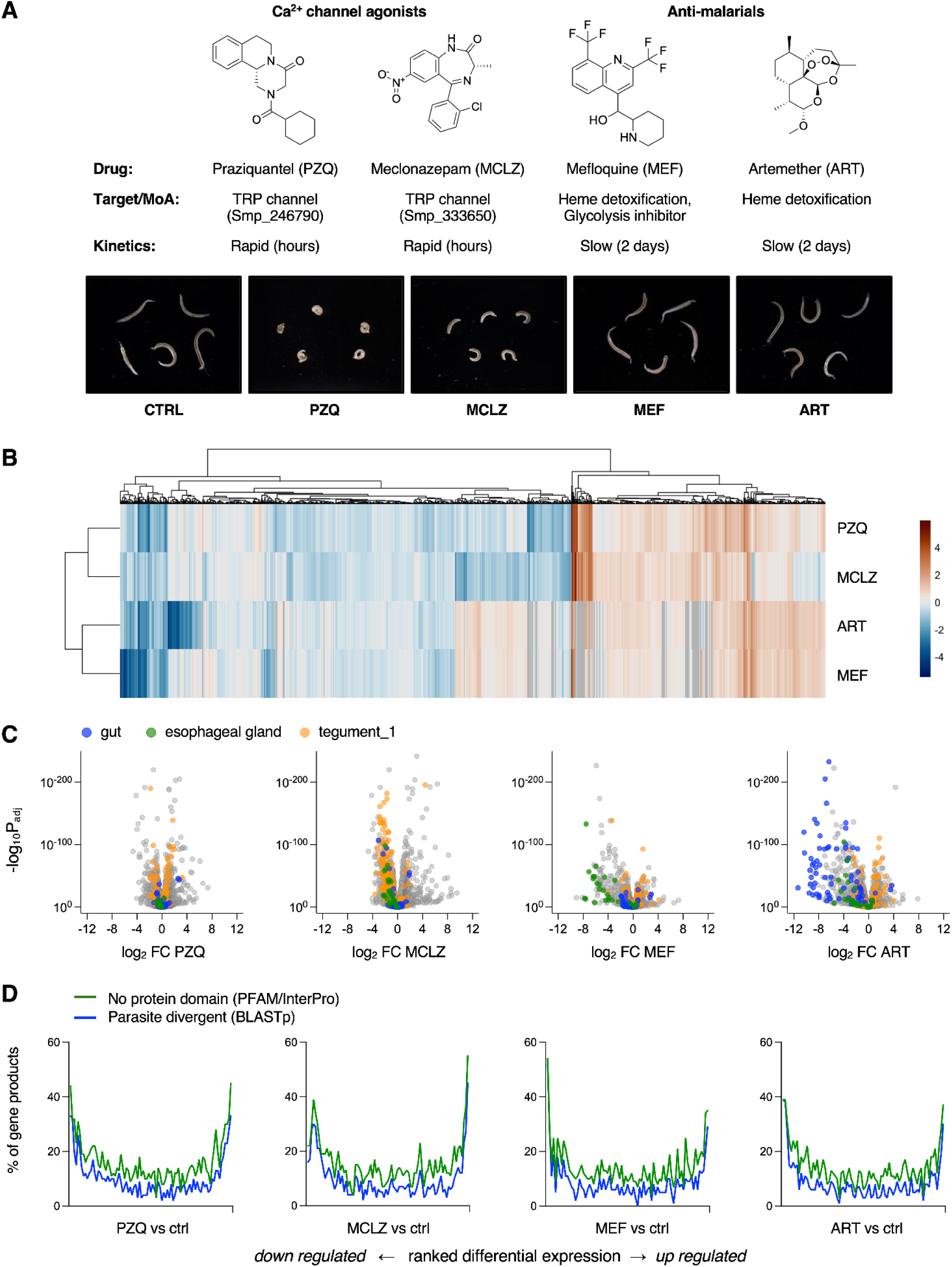
Comparative effects of ion channel agonists and antimalarial drugs on adult *S. mansoni.* **(A)** Top -summary of TRP agonist and antimalarial chemical structures and putative mechanisms of action. Bottom -images of worms treated in vitro with meclonazepam 5 µM (MCLZ), praziquantel 1 µM (PZQ), mefloquine 50 µM (MEF), or artemether 50 µM (ART). **(B)** Hierarchical clustering of gene products differentially expressed in at least one treatment condition (1,588 genes; 74 DEGs were detected in only one treatment cohort and are not shown), with TRP ligands (MCLZ and PZQ) and antimalarials (ART and MEF) clustering together. Blue = down-regulated transcripts. Brown = up-regulated transcripts. Scale = log_2_ fold change. **(C)** Volcano plots of bulk transcriptomic response of adult worms to in vivo treatment with each of the above compounds (PZQ 100 mg/kg, MCLZ 30 mg/kg, MEF 200 mg/kg, ART 300 mg/kg). Colors denote changes to cell and tissue-specific markers identified in prior single-cell transcriptomic work (32); orange = tegument_1 cluster, green = esophageal gland cluster, blue = gut cluster. **(D)** In all treatment conditions, differentially expressed transcripts tended to be enriched in unannotated or parasite-specific gene products. Transcripts are arranged in rank order from most down-regulated (left side of the axis) to most up-regulated (right side of the axis). For each window of 100 genes, the percentage with no annotated protein domain (PFAM or InterPro) or no BLASTp hit in free-living flatworms is shown.

Mice harboring patent infections with sexually mature parasites (n = 5 mice per group) were dosed with MEF (200 mg/kg) and ART (300 mg/kg), and worms were collected from mouse livers after 48 hours. Bulk RNA-Seq was performed on these samples, and reads were mapped to version 10 of the *S. mansoni* reference genome (30, 31). Principal component analysis showed clear separation of control, MEF and ART treated cohorts (**Supplemental Figure 1**). Differentially expressed genes (DEGs) were identified using thresholds of absolute log_2_ fold change >1 and a false discovery rate (FDR) adjusted p value < 0.01. These results were compared with published datasets from PZQ-treated worms (25), which we previously had mapped to version 7 of the *S. mansoni* genome and have remapped here to version 10, and MCLZ treatment from previously published work (26). All of these datasets were generated in the same murine model (female Swiss Webster mice), using the same *S. mansoni* strain (adult NMRI), and processed by the same bioinformatic pipelines. In total, this resulted in 552 DEGs following PZQ treatment, 803 DEGs following MCLZ treatment, 937 DEGs following ART treatment, and 460 DEGs following MEF treatment (**Supplemental Files 1 & 2**). Hierarchical clustering of the union of these datasets (1662 unique gene products) grouped the two TRP agonist treatments (PZQ and MCLZ) separately from the two antimalarial treatments (ART and MEF) (**Figure 1B**). However, each treatment evoked largely distinct changes in gene expression. Only 33 transcripts were up-regulated and 63 transcripts were down-regulated across all four treatment datasets. These distinctions are also apparent when examining changes in markers of specific cell and tissue types defined by a published adult *S. mansoni* single cell RNA-Seq atlas (32) (**Supplemental Table 1**). For example, while all treatments altered gene expression in the tegument, ART caused a pronounced down-regulation of gut / intestinal cell markers (**Figure 1C**). Finally, transcriptional changes tended to be most pronounced for genes that are more divergent in parasitic flatworms (**Figure 1D**). We previously noted this profile with TRP agonists PZQ (25) and MCLZ (26), where up- and down-regulated transcripts were enriched for gene products with no discernable orthologs outside of parasitic flatworms. We assessed this pattern using the absence of annotated protein domains (Pfam or InterPro, **Supplemental File 3**) and the absence of detectable BLASTp homologs in free-living members of the Platyhelminth phylum (26). While this trend was clear for each drug treatment, the exact gene products tended to be unique for each dataset, consistent with the limited overlap in up- and down-regulated DEG lists noted above.

### Transcriptional response to TRP channel agonists

PZQ and MCLZ have very similar phenotypes (contractile paralysis, tegument damage) but activate different schistosome TRP channels. Transcriptomic data indicate that 50.4% of PZQ-responsive DEGs and 34.6% of MCLZ-responsive DEGs are shared and change in the same direction (**Figure 2A**). When comparing these DEG lists, we found that many of the most strongly up-regulated or down-regulated genes were unannotated in the most recent version of the *S. mansoni* genome on WormBase ParaSite (WBP19). For example, 33 of the top 100 transcripts changing with MCLZ treatment and 17 of the top 100 transcripts changing with PZQ treatment are unannotated. Visualizing the gene products up- and down-regulated with both treatments in a sequence similarity network shows that many of the unannotated gene products are similar to each other (**Figure 2B**). Members of this cluster belong to a recently reported subtelomeric gene family characterized by the presence of Plexin-Semaphorin-Integrin (PSI) domains (33). These genes are primarily expressed in the tegument of the intra-molluscan parasite life-cycle stages (33, 34), and were originally termed ‘CP391S-like egg proteins’ in early EST databases (35). Single members of the family are present in other trematodes such as the liver fluke *Clonorchis sinensis* and the avian schistosome *Trichobilharzia szidati*, but the family appears to have expanded in both Asian and African schistosome species (**Figure 2C, Supplemental File 4**). Available gene expression data across life cycle stages confirms high expression during intra-molluscan stages and low expression during intra-mammalian stages (36, 37) (**Figure 2C**). However, the transcriptional change in this gene family was very similar in both PZQ and MCLZ treated adult parasites, with similar degrees of up-regulation relative to control samples following drug exposure (**Figure 2D**). Because of the location of this gene family near the ends of chromosomes, many of these genes were not present in older genome assemblies, and prior work on PZQ-evoked changes in gene expression (including our earlier RNA-Seq study) did not note the extent of the transcriptional changes in this family following drug treatment. Version 10 of the *S. mansoni* genome has greatly improved coverage of these sub-telomeric regions. Reanalysis of our earlier dataset (25) and older studies on PZQ-evoked transcriptional changes (PRJEB1281), both remapped to the *S. mansoni* v10 genome, showed that members of this family are up-regulated following PZQ exposure. This observation is replicated across multiple datasets from different laboratories and different strains of *S. mansoni* (**Figure 2E**).

**Figure 2.**
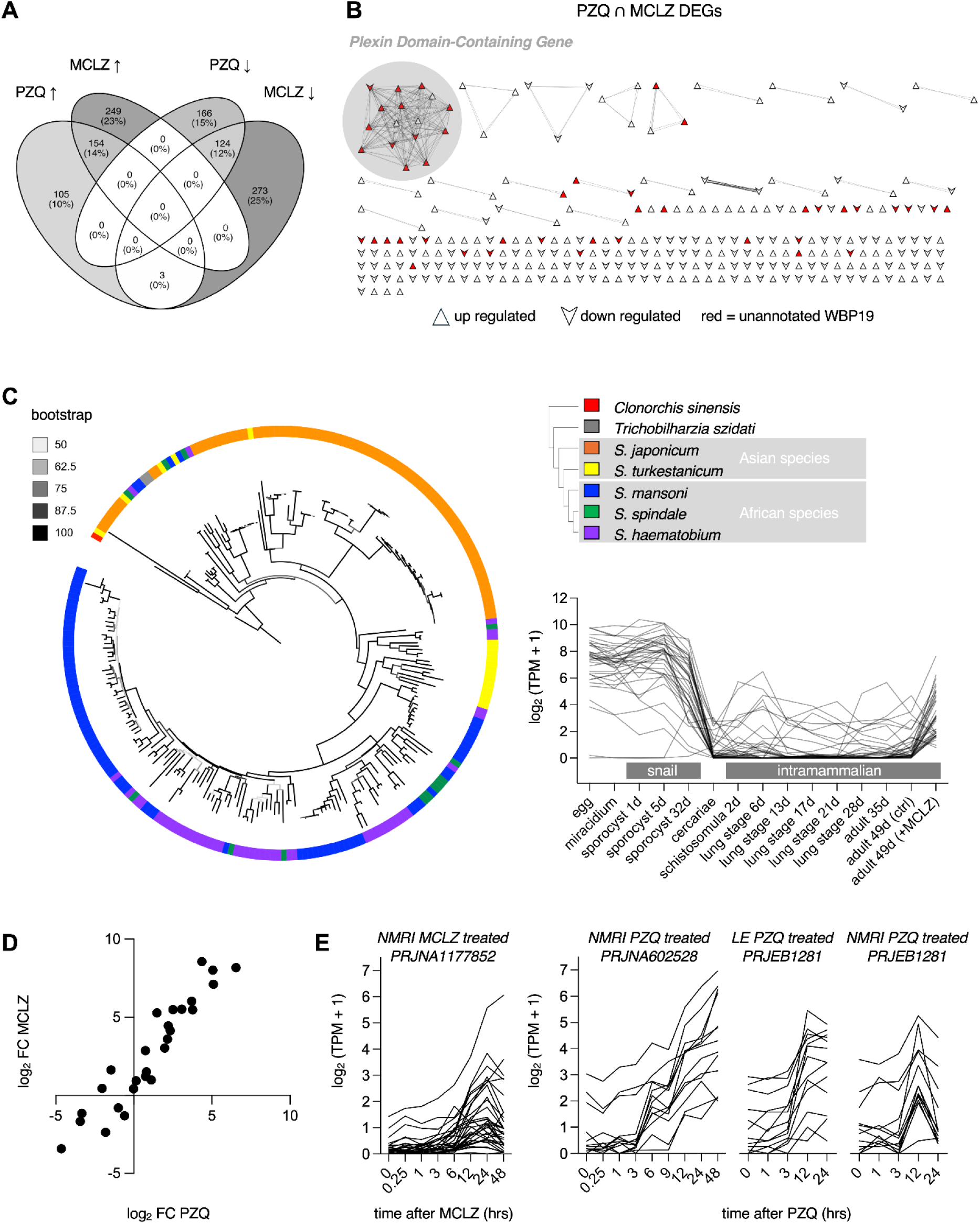
TRP ligands meclonazepam and praziquantel increase expression of plexin-domain containing genes. **(A)** Venn diagram of differentially expressed transcripts in *S. mansoni* following MCLZ or PZQ exposure. **(B)** Sequence similarity network of gene products differentially expressed with both treatments. Many are unannotated in the most recent version of the *S. mansoni* genome, but cluster together and belong to the plexin domain-containing gene family. Red symbols = genes with no annotation in WBP19. Upward triangle = up-regulated transcripts. Downward triangle = down-regulated transcripts. **(C)** Phylogenetic tree of plexin domain-containing gene products, revealing limited orthologs in other trematodes (e.g., liver fluke *Clonorchis sinensis*, avian blood fluke *Trichobilharzia szidati*) but expansion in both African and Asian *Schistosoma* species. Inset -gene expression data across the *S. mansoni* life cycle for the 16 plexin domain-containing genes highlighted in B, showing higher expression in larval, intramolluscan stages. Expression in log_2_ (TPM +1) is shown using data from transcriptomic datasets for egg to schistosomula stages (PRJEB32839, (37)), lung stages to adult 35 d (PRJEB30644, (36)), and adult 49 d (7 weeks) with and without MCLZ treatment (PRJNA1150591, (26)). **(D)** Correlation of changes in expression for the plexin domain-containing genes in adult *S. mansoni* following treatment with either PZQ (remapped data from reference (25); PRJNA597909) or MCLZ (data from reference (26); PRJNA1150591), relative to control. **(E)** Time course of gene expression changes following drug treatment; (i) MCLZ treatment (PRJNA1177852), (ii) PZQ treatment (PRJNA602528), (iii & iv) PZQ treatment (data from PRJEB1281, WBP19).

### Transcriptional response to antimalarial treatments

Antimalarial drugs ART and MEF have demonstrated antischistosomal efficacy in both murine models (18–20) and human studies (21, 38–40), but less work has been done profiling the parasite response to these drugs at the molecular level. The transcriptional response to these two drugs was more similar to each other than to either of the TRP agonists, with ART treatment resulting in a greater number of DEGs than MEF. However, many genes differentially expressed with MEF treatment were also present in the ART dataset; 85% of transcripts up-regulated with MEF were also up-regulated with ART, and 91% of transcripts down-regulated with MEF were also down-regulated with ART (**Figure 3A**). Sequence similarity networks were again used to assess the types of gene products changing with each drug treatment, clustering the top 100 up- and down-regulated gene products (ranked by log_2_ fold change) for each condition (**Figure 3B**). The networks helped identify related genes that were not obvious from electronic gene annotations, because some gene products with high sequence similarity have different annotations. The two treatments, ART and MEF, were similar in terms of up-regulated gene products, with plexin domain-containing genes again represented. Greater differences were observed in down-regulated gene products, with ART treatment down-regulating gene families with roles in feeding and digestion (**Figure 3C**). Therefore, the two treatments were similar in terms of the most up-regulated gene products, but showed greater differences in the most down-regulated gene products (**Figure 3D**).

**Figure 3.**
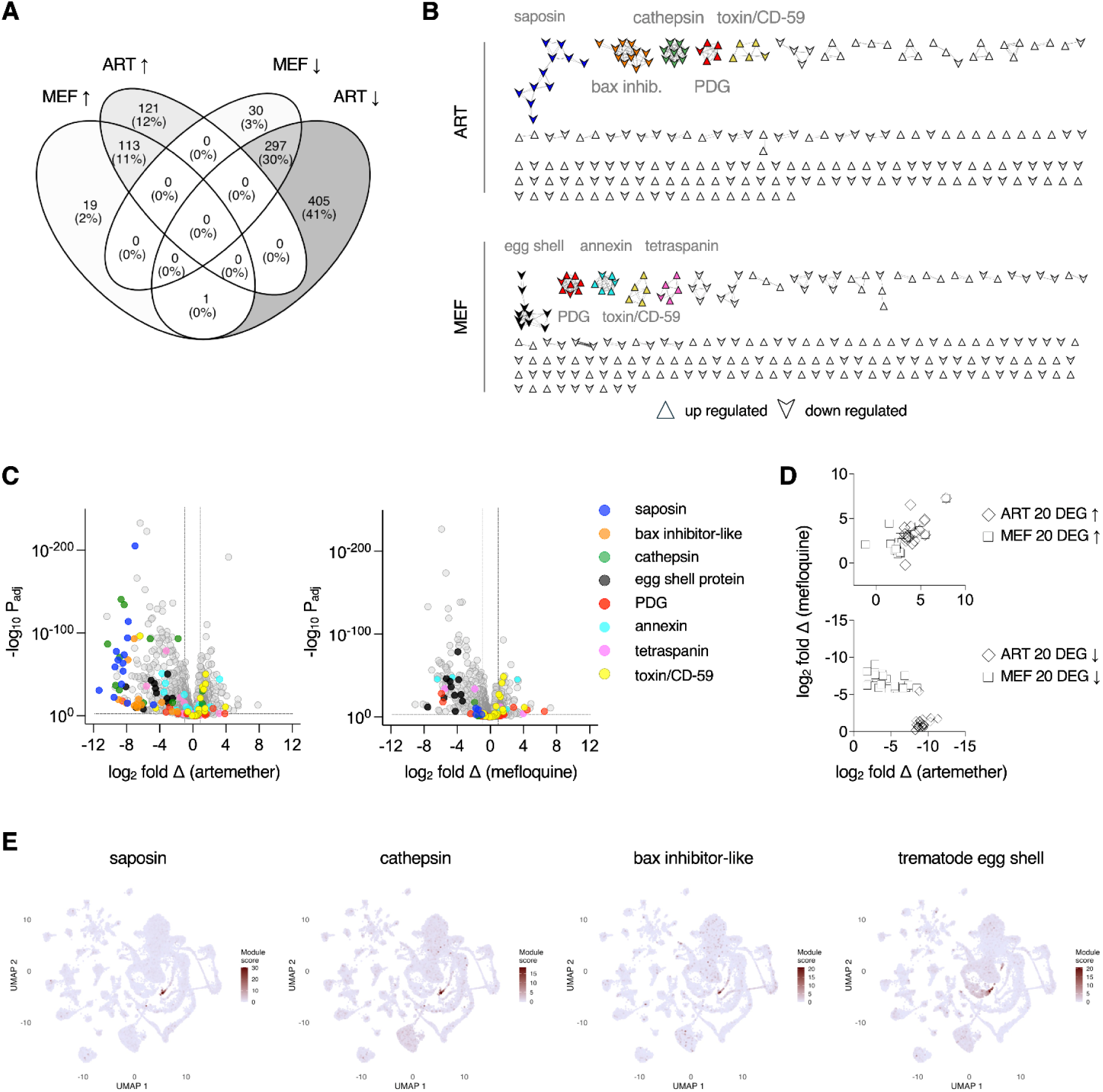
Transcriptional response to antimalarial drugs artemether and mefloquine. **(A)** Venn diagram of differentially expressed transcripts in *S. mansoni* following ART or MEF exposure. **(B)** Sequence similarity network of the top 100 up- and down-regulated gene products for each treatment. The top five clusters for each treatment are colored indicating the following gene families; saposins (blue), bax inhibitor-like (orange), cathepsin (green), plexin domain-containing gene (PDG, red), CD59-like (yellow), egg shell protein (black), annexin (cyan), and tetraspanin (pink). **(C)** Volcano plot of gene expression differences between ART (left) and MEF (right), relative to control. Selected gene families are shown colored as in B. **(D)** Correlation of changes in gene expression in ART and MEF treated samples. The top 20 up- and down-regulated gene products are shown for ART treated samples (diamond) and MEF treated samples (squares), relative to controls. **(E)** Module scores for genes with characteristic protein domains that are differentially expressed (cutoff of log_2_ fold change >1 and adjusted p value < 0.01) following ART (saposin, cathepsin and bax inhibitor) or MEF (trematode egg shell) treatment.

To examine the cellular expression of the gene families that showed prominent changes with ART or MEF, we mapped the differentially expressed members of the saposin, cathepsin, Bax inhibitor-like, and trematode eggshell protein families onto the adult *S. mansoni* single cell gene expression atlas (32). For each gene family, we standardized the expression of each DEG across cells (converting to a z-score), averaged these values within each cell to generate a gene-set expression score, and visualized these on the published UMAP embedding (**Figure 3E**). The ART-responsive saposin, cathepsin and Bax-inhibitor-like families showed prominent expression within clusters annotated as gut cells. This localization is consistent with a potential role for heme-dependent activation of artemether, as described for artemisinin-derivatives in malaria parasites (41–43). On the other hand, pronounced changes to these gut markers were not observed with MEF treatment. Instead, gene products related to eggshell formation were among the most strongly down-regulated with MEF. These were also down-regulated with ART, but ranked lower given more extreme changes in groups such as saposin genes. Because eggshell proteins are made in vitellaria, these changes may reflect the effects of antimalarial treatment action on female reproductive tissues (**Figure 3B,C&E**). Decreased expression of these vitellocyte markers does not appear to reflect a bias in sample composition (a lower proportion of female worms), because the median log_2_ fold change of testis and ovary markers was close to zero for all four drug treatments (testis −0.01 to +0.10; ovary −0.22 to −0.06; **Supplemental Figure 2**).

### In vivo efficacy of praziquantel-alternatives in a drug-resistant model of parasite infection

A laboratory-selected, PZQ-resistant *S. mansoni* population (SmLE-PZQ-ER) has been shown to be more than 300-fold less responsive to PZQ in vitro than the susceptible population (SmLE-PZQ-ES) (5). We used these populations to test for cross-resistance between PZQ and potential alternatives (MCLZ, MEF, and ART). Cross-resistance may occur when drugs share a target or resistance mechanism. Although these compounds likely act through different targets, each has been linked to Ca^2+^ dysregulation through various mechanisms (44–46). Artemisinin derivatives have also been directly implicated as TRP agonists (47, 48), although we did not observe activity at schistosome TRP_PZQ_ or TRP_MCLZ_ channels (**Supplemental Figure 3**).

To test for cross-resistance, we dosed mice infected with adult SmLE-PZQ-ES or SmLE-PZQ-ER parasites once orally with PZQ (200 or 400 mg/kg), MCLZ (30 mg/kg), ART (300 mg/kg) or MEF (200 mg/kg) and recovered surviving worms one week later. We selected doses near the lower end of published effective ranges to improve sensitivity to differences in drug efficacy between parasite populations (**Figure 4, Supplemental Table 2**). At the 400 mg/kg dose, PZQ eliminated all sensitive worms and reduced resistant male and female worm burdens by 78.6% and 82.9%. The difference between sensitive and resistant lines was significant in males (p = 0.008) and nominal in females (P = 0.048). PZQ 200 mg/kg reduced sensitive male worms by 99.0%, but resistant male worms by only 39.3% with no significant difference from control worm burden (P = 0.33). In contrast, MEF and ART reduced worm burden by at least 97% in both sexes of both populations. MCLZ reduced worm burden by 90–92% in sensitive and 80–83% in resistant parasites. None of these drugs differed significantly between populations (p ≥ 0.24).

**Figure 4.**
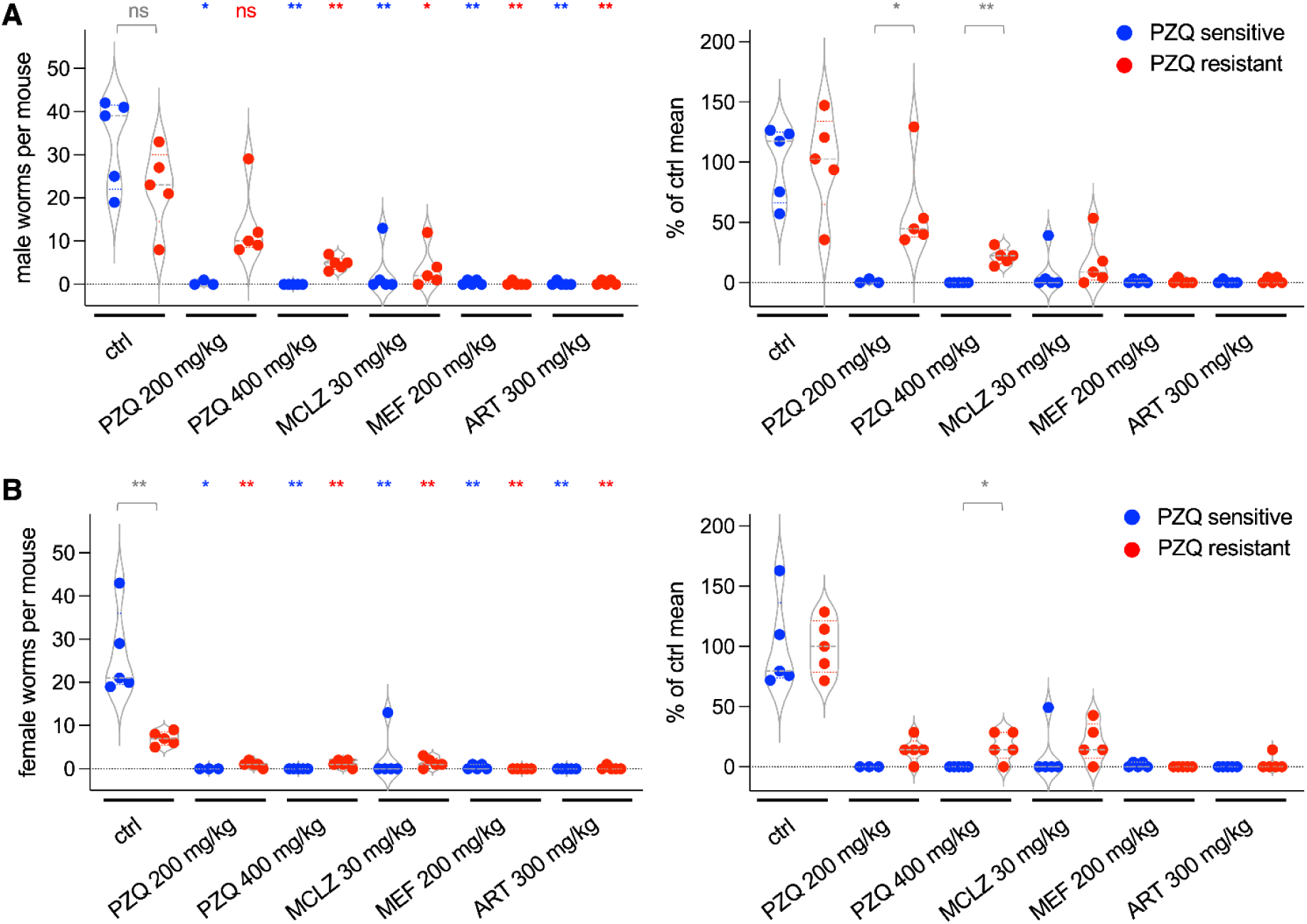
In vivo efficacy of various drug classes in praziquantel sensitive and resistant populations of *S. mansoni*. Mice harboring adult PZQ-sensitive (SmLE-PZQ-ES, blue) or PZQ-resistant (SmLE-PZQ-ER, red) *S. mansoni* were dosed once orally with PZQ (200 or 400 mg/kg), MCLZ (30 mg/kg), MEF (200 mg/kg) or ART (300 mg/kg) and adult worms were recovered one week later. Counts are shown of **(A)** male and **(B)** female worms, with number of worms recovered from each mouse shown on the left and the same worm burden normalized as percentage of the mean of the control group of the same parasite population (sensitive or resistant) on the right. Each data point reflects the worm burden of one mouse, n = 5 per group, with the exception of PZQ-sensitive, PZQ 200 mg/kg (n = 3), where two mice in the cohort were euthanised before treatment began. Males and females were analysed separately with two-sided exact Wilcoxon–Mann–Whitney tests. * p < 0.05; ** p < 0.01; ns = not significant. P values and Holm-adjusted values for multiple comparisons are provided in Supplemental Table 2. Left -asterisks above the blue and red dots compare each drug group with the ctrl group of the same line (sensitive or resistant). Grey bracket compares sensitive and resistant parasite burdens in control mice. Right -grey brackets compare sensitive and resistant parasite burdens for each drug; comparisons without a bracket were not significant.

## Discussion

Reliance on praziquantel monotherapy for schistosomiasis raises concerns of selection for drug-resistant parasites and suboptimal control of the disease due to lack of efficacy against juvenile worms (49, 50). These concerns may be addressed by incorporating drugs with alternative mechanisms of action. These classes such as antimalarials (**Figure 5A**), which are also highly effective against schistosomes (14, 51–53), or compounds that act on other ion channels (**Figure 5B**). Here, we present a detailed study of several PZQ-alternatives using comparative RNA-Seq to identify similarities and differences between the various chemotherapies.

**Figure 5.**
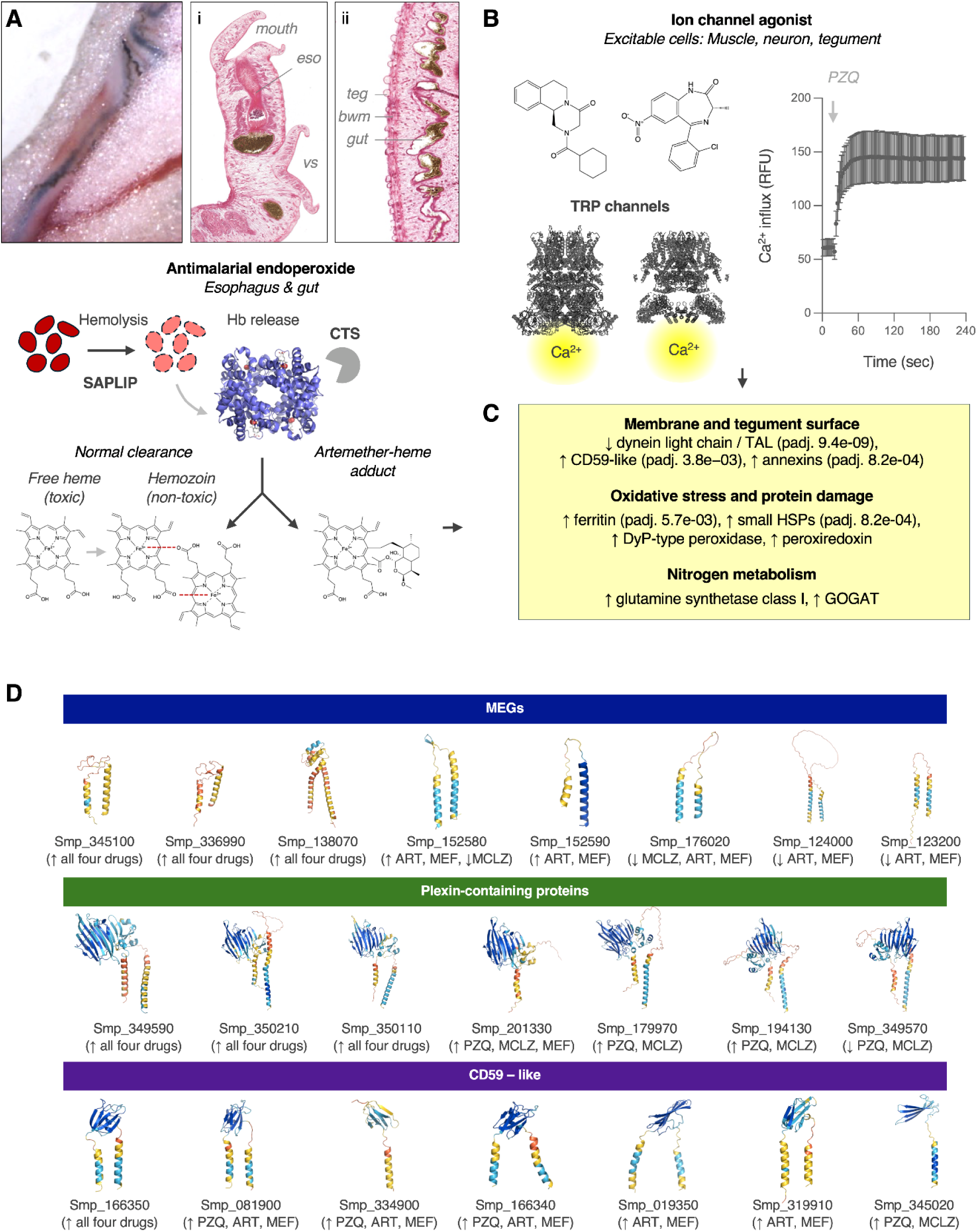
Action of antimalarials and TRP agonists on *S. mansoni*. **(A)** Images of adult schistosomes showing tissues with high heme and iron content. Left -image of paired male and female worms within the mesenteric vasculature. Dark gut contents and female vitelleria are visible. Right -H&E stained sections of the anterior and dorsal regions of an adult male parasite, showing esophageal (eso) and intestinal contents from blood feeding (dark brown). vs = ventral sucker, teg = tegument, bwm = body wall muscle. These may reflect sites of action for ART, given the transcriptional response showing changes in gene products involved in hemolysis (saposin-like proteins, SAPLIP) and hemoglobin (Hb) digestion (CTS, cathepsin). Under normal conditions, heme released by Hb digestion polymerizes into hemozoin. ART may impair this process. **(B)** Actions of TRP agonists. Structures of PZQ (left) and MCLZ (right), above homology models of their respective targets; Smp_246790 (AlphaFold DB AF-A0A5K4F442-F1) superposed onto PDB 6PUS, and Smp_333650 (AlphaFold DB AF-A0A5K4FCC0-F1) superposed on PDB 5VKQ with MODELLER 10.8. Regions with pLDDT < 50 not shown. **(C)** Treatment with both TRP and antimalarial drug classes results in changes to gene products expressed at the host-parasite interface, as well as pathways involved in oxidative stress and metabolism. Adjusted P values (Fisher exact test with Benjamini-Hochberg correction) reflect enrichment of InterPro domains. Domains present in four or fewer proteins, or with only one member in the DEG set, were not tested for enrichment. **(D)** However, enrichment analysis can miss some families of parasite-specific esophageal gland or tegument-expressed small secreted and surface proteins. In addition to the CD59-like genes found by InterPro analysis, micro-exon and plexin-containing genes are also enriched (padj. 2.0e−12 and 2.9e−10, respectively).

### Artemether action on gut cells

Despite evidence from human and animal studies that antimalarial chemotherapies are effective at treating schistosomiasis, there have been few studies into their antiparasitic molecular mechanism of action in flatworms. One key finding is that artemisinin derivatives are not active against schistosomes in vitro unless the culture media is supplemented with haemin (54, 55). In *Plasmodium*, ferrous heme is required for cleavage of the endoperoxide bridge of artemisinin derivatives and generation of reactive intermediates (42, 56) that can modify biological macromolecules (43). This can include formation of heme-drug adducts that may inhibit hemozoin formation (57–59). Covalent modification of cellular proteins or nucleic acids or interference with hemozoin formation and heme detoxification may also be effective antischistosomal mechanisms. Our RNA-Seq data identify a pronounced change in transcriptional markers of intestinal cells (**Figure 3, Supplemental Table 1**), consistent with previous ultrastructural studies reporting artemether-induced damage to these tissues in *S. mansoni* (60). The schistosome gut is also where heme and hemozoin are concentrated following ingestion of red blood cells (**Figure 5A**), and the transcriptional signature observed for these markers with artemether, but not the three other drugs, is consistent with a heme or hemozoin dependant mechanism in vivo. This includes down-regulation of saposin-like proteins, which are involved in lysis of ingested blood cells across a range of species of hematophagous parasitic worms (61–65), as well as cathepsins, which degrade and digest hemoglobin and are required for parasite growth and development (66–70).

### Transcriptional responses observed in response to both drug classes

While artemether treatment uniquely impacted gut cells, certain transcriptional changes were observed across drug classes. These include various families of small secreted proteins (e.g. micro-exon genes (MEGs), plexin domain containing proteins, and CD59 like proteins), as well as gene products involved in redox dysfunction (**Figure 5C&D**). These changes may reflect shared parasite stressors such as impaired feeding, tegment damage, or a response to attack by host immune cells.

Esophageal markers were differentially expressed with the TRP agonist MCLZ and antimalarials MEF and ART (**Supplemental Table 1**). The lack of signal in the PZQ dataset may be due to the dose and timepoint (100 mg/kg, 14 hours) in that experiment, since changes in these markers emerge later in the time course with 400 mg/kg treatment (25). The esophageal gland processes ingested blood cells (71, 72) and contributes to host immune evasion (73), with secreted gene products such as MEGs (74) and Kunitz-type protease inhibitors (75) involved in lysis of blood cells and anticoagulation. Members of these families, including MEG-2 and MEG-3, were up-regulated in prior RNA-Seq analysis of *S. mansoni* responses to PZQ and MCLZ (25, 26), and are also up-regulated with MEF and ART (MEGs Smp_138070, Smp_336990 and Smp_345100, and the Kunitz-type protease inhibitor Smp_311670). Earlier work proposed these changes reflect a response to TRP agonists (25, 26), but their induction following antimalarial treatment suggests a broader response to drug-induced damage that may include evasion of host immune attack.

The requirement for immune evasion may be driven by tegument damage caused by each drug treatment. Many of the gene products up-regulated across different drug treatments have little or no expression in adults, but are highly expressed during larval life cycle stages. The reexpression of larval stage genes in treated adults may reflect immune pressures shared by newly invading and drug-damaged parasites. Penetrating cercariae and schistosomules are susceptible to host complement attack immediately following infection. As they mature, schistosomules acquire protection with the development of a heptalaminate tegument membrane which absorbs host derived molecules (76–78). Drug treatment damages the adult tegument and renders the parasite vulnerable to host immune attack (25, 26, 79–81). This may be why larval gene products are re-expressed. Examples include the secreted helminth defence molecule Sm16 (Smp_341790), which is highly expressed in invading cercariae and inhibits macrophage activation (82–84). The MEGs noted above also fit this trend, as do plexin domain-containing genes, which are highly expressed in the developing tegument in earlier life cycle stages (33, 34, 85). Given their localization at the tegument and sub-telomeric placement in the genome, plexin domain-containing genes have been suggested to have roles in immune evasion similar to *Trypanosoma* VSG and *Plasmodium* var genes genes (33). Alternatively, reactivation of genes expressed during tegument development may also reflect a response aimed at repairing damage to this tissue following chemotherapy (86).

### Prospective targets and pathways for drug development

Several pathways highlighted by these data may offer opportunities for anthelmintic drug development because they contain proteins absent from vertebrate hosts or exploit vulnerabilities associated with the schistosomes blood-feeding lifestyle and exposure to large amounts of iron and hemoproteins.

The most strongly up-regulated gene in PZQ, ART and MEF treated samples was Smp_133770, a putative class I glutamine synthetase (**Supplemental Figure 4**). (Smp_133770 was also up-regulated with MCLZ, but the adjusted p value = 0.02 was above our selected threshold of 0.01). This is unusual, because class I glutamine synthetases are typically regarded as prokaryotic enzymes (87, 88). Class I glutamine synthetase homologs also occur in some eukaryotes. Most of these, such as the vertebrate lens protein lengsin, are catalytically inactive and have evolved structural roles (89). A catalytically active class I glutamine synthetase has been reported in *Plasmodium* parasites, which also digest host hemoglobin and metabolize the resulting amino acids. To our knowledge, this is the only eukaryotic example of a catalytically active class I glutamine synthetase (90, 91). Smp_133770 may have a structural role in response to drug-induced damage, but the protein may also be catalytically active, as it retains residues conserved in active prokaryotic and *Plasmodium* orthologs but lost in inactive animal lengsins (89, 92). The glutamine cycle has important roles in nitrogen metabolism and ammonia detoxification in protozoan parasites (93), molting and development in nematode parasites (94), and maintenance of tegument membrane potential in schistosomes (95). In addition to glutamine synthetase, invertebrates possess glutamate synthase (GOGAT; Smp_128370, up-regulated in PZQ, MEF and ART treated samples), which transfers the glutamine amide nitrogen to 2-oxoglutarate, producing two glutamate molecules. Together, glutamine synthetase and GOGAT mediate the incorporation of free ammonia into glutamate. Because vertebrates lack GOGAT, and human glutamine synthetase is class II rather than class I, both of these enzymes may be parasite-selective drug targets.

Genes encoding proteins potentially involved in iron-homeostasis and oxidative damage were also up-regulated following all four treatment conditions. Peroxide in the presence of free iron (Fe^2+^) is particularly damaging because it generates highly reactive hydroxyl radicals. A protective response could conceivably lower peroxide levels and sequester free iron. The heme-containing peroxidase Smp_314210 is a DyP-type enzyme, a family not present in known mammalian genomes. DyP-type peroxidases are sparsely distributed in metazoan genomes (96–98). The only biochemically characterized animal DyP-type peroxidase is from the flatworm *Echinococcus multilocularis* (99), where it was experimentally shown to exhibit catalase activity. The possibility that Smp_314210 may also exhibit this property is notable, because the schistosome genome otherwise lacks a predicted catalase enzyme (100) and no catalase domain (Pfam PF00199) is annotated in the *S. mansoni* proteome (**Supplemental File 3**). Free iron may also be sequestered by ferritin, while simultaneously lowering peroxide levels. Ferritins sequester iron within their core, in the process oxidising Fe²⁺ to Fe³⁺ while reducing H₂O₂ to water. Ferritin paralogs are up-regulated in all four treatments (Smp_311630, 3.8 to 13-fold, rising from 48–79 to 316–1,054 TPM; Smp_311640, 3.6 to 11.6-fold, 16–28 to 89–342 TPM).

## Conclusions

Comparative RNA-Seq showed transcriptional profiles clustered by drug class, providing insight into the distinct mechanisms of these antischistosomal chemotherapies. However, numerous gene families also exhibited shared changes across experimental treatments, perhaps reflecting parasite responses to general redox dysfunction or attack by the host immune system that may also be exploited for anthelmintic development. We further show that MEF, ART and MCLZ retain activity against a laboratory-selected praziquantel-resistant population, consistent with the different targets of these compounds and supporting their further investigation as alternatives to praziquantel monotherapy.

## Materials and Methods

### Ethics statement

All animal experiments were conducted in accordance with the Animal Welfare Act and the Health Research Extension Act. Experiments were approved by the Medical College of Wisconsin Institutional Animal Care and Use Committee (IACUC) protocols AUA00006471 and AUA00006735 and UW-Madison School of Veterinary Medicine IACUC (protocol V006353).

### Parasite drug treatment assays

Female Swiss Webster mice infected with the NMRI population of *S. mansoni* (∼180 cercariae per mouse) were obtained from the NIH-NIAID Schistosomiasis Resource Center (SRC). Drugs were obtained from the following vendors; praziquantel (Enzo Life Sciences), meclonazepam (Anant Pharmaceuticals), artemether (Tokyo Chemical Industry America), mefloquine (Sigma-Aldrich). For in vitro drug assays, mice were euthanized by CO_2_ asphyxiation and cervical dislocation seven weeks post-infection. Adult schistosomes were dissected from the mesenteric vasculature and cultured in high-glucose DMEM supplemented with 5% fetal calf serum, penicillin-streptomycin (100 units/mL), HEPES (25 mM) and sodium pyruvate (1 mM) at 37°C in 5% CO_2_ supplemented with either anti-parasitic compound or DMSO vehicle control for 24 hours, and images of drug treated worms were acquired with a stereomicroscope. For in vivo drug treatments prior to RNA-Seq studies, mice harboring patent seven week old *S. mansoni* infections were treated with either DMSO vehicle control, artemether (ART, 300 mg/kg), or mefloquine (MEF, 200 mg/kg) solubilized in vegetable oil (0.25 mL) by oral gavage (n=5 mice per group). These doses were selected as they are within the range established by prior literature using a murine model of *S. mansoni* infection (18–20). 48 hours after treatment, mice were euthanized and worms were recovered from the liver, homogenized in TRIzol reagent (Invitrogen), and stored at -80°C until processing for RNA-Seq. For experiments comparing the efficacy of test compounds on PZQ sensitive and resistant schistosome populations, female Swiss Webster mice infected with SmLE-PZQ-ER (Enriched Resistant) and SmLE-PZQ-ES (Enriched-Sensitive) parasite populations described in reference (5) were obtained from the NIH-NIAID SRC. At seven weeks postinfection, mice were treated with PZQ, MCLZ, ART and MEF as outlined above, solubilizing compounds in vegetable oil and administering by oral gavage. Drug efficacy was assessed one week later, euthanizing mice and dissecting parasites from the mesenteric vasculature and liver to count surviving worms. Worm burdens were compared between treated and vehicle control mice within each parasite population (SmLE-PZQ-ES and SmLE-PZQ-ER), and percentage worm burden reductions were compared between the two populations, using two-sided exact Wilcoxon-Mann-Whitney tests. Unadjusted and Holm-adjusted P values are reported in **Supplementary Table 2**. P < 0.05 was considered significant.

### Transcriptome sequencing

Total RNA was extracted from parasites dissected from DMSO control, MEF and ART treated mice using TRIzol reagent (Invitrogen), generating one library per mouse. Libraries were prepared using the TruSeq Stranded mRNA kit (Illumina) and sequenced on an Illumina HiSeq 2500 (50 bp paired-end reads, 11–17 million reads per sample). RNA-Seq data from control, ART-treated and MEF-treated samples have been deposited to NCBI (controls SAMN64214095 -SAMN64214099, artemether SAMN64214090 -SAMN64214094, and mefloquine SAMN64214100 -SAMN64214104). Gene descriptions and protein domain annotations were obtained from WormBase ParaSite. In addition to the antimalarial treatments ART and MEF, previously deposited samples from PZQ treated worms (SAMN13691171 -80) from reference (25) were re-mapped, and data from MCLZ treatment was retrieved from reference (26). The RNA-Seq analysis pipeline was implemented using Nextflow (101) and is available at (https://github.com/zamanianlab/Core_RNAseq-nf). Reads were trimmed using fastp (v0.20.1) (102) and aligned using STAR (v2.7.6a) (103) to the *Schistosoma mansoni* reference genome (version 10) (31) which was obtained from Wormbase Parasite release 19 (30). Overlapping mates were merged when they overlapped by at least 10 bases. The proportion of read pairs mapping uniquely was 70.8 -83.9% for the control, ART and MEF libraries, 86.3 -88.5% for the PZQ libraries and 65.1 -79.6% for the MCLZ libraries.

### Differential expression analysis

Differential expression was analyzed separately for each experiment with DESeq2 (v1.50.2) (104). The PZQ and MCLZ experiments were analyzed as two-group comparisons (treatment vs control). The ART and MEF libraries were generated in a single experiment with one shared control group, and were analyzed in one three-group model (control, ART and MEF). Genes with at least 10 reads in at least five libraries were retained, and results were obtained with default settings (Wald test, Benjamini-Hochberg adjustment). Differentially expressed genes (DEGs) were defined as FDR adjusted P value < 0.01 and absolute log_2_ fold change > 1, using unshrunken fold change estimates. Principal component analysis was performed on the control, ART and MEF experiment (**Supplemental Figure 1**), transforming counts with the DESeq2 variance-stabilizing transformation and using the 500 genes with the highest variance across libraries. Principal components were calculated on the centred, unscaled transformed values with the DESeq2 plotPCA function.

To determine whether the observed changes in female-enriched transcripts (vitellocyte markers) were due to differences in the proportion of female worms between samples, we assessed the expression of germline markers from the adult *S. mansoni* single-cell atlas (32). Log_2_ fold change values markers in testis (late male germ cells) and ovary (late female germ cell) clusters were used to calculate the median change for each marker set (**Supplemental Figure 2**).

### Bioinformatic analysis of gene families

Sequence similarity networks were generated for lists of genes by performing an all-versus-all BLASTp search of the protein sequences under investigation, with the resulting bit scores used to weight interactions in the network. Networks were visualized in Cytoscape (v3.10.4). Plexin domain-containing genes were analyzed using the *S. mansoni* sequences in reference (33) to perform a BLASTp search against the predicted proteomes of the following parasitic flatworms; *Clonorchis sinensis* (PRJNA386618), *Trichobilharzia szidati* (PRJEB44434), *S. japonicum* (PRJNA739049), *S. turkestanicum* (PRJEB44434), *S. mansoni* (PRJEA36577), *S. spindale* (PRJEB44434) *and S. haematobium* (PRJNA78265). Reciprocal BLASTp searches against *S. mansoni* were then performed, retaining sequences whose top hits matched one of the original plexin domain-containing proteins from reference (33). These candidates were then aligned using Clustal Omega (105) and trimmed with TrimAl (v1.5, (106)). Sequences with conserved cysteines in PSI domain (specifically, the CxWC motif) were used to generate a phylogenetic tree with IQ-TREE (v2, (107)) with 1000 bootstrap replicates, which was then visualized in iTOL (v7, (108)). Publicly available gene expression data for plexin domain-containing genes were retrieved from WormBase Parasite; expression across egg and intra-molluscan life cycle stages was retrieved from (37) (European Nucleotide Archive accession number PRJEB32839) and expression across intramammalian life cycle stages was retrieved from (36) (ERP113121). These RNA-Seq datasets were selected for their temporal resolution of different life cycle stages. As neither of these studies include the 49 day-post-infection timepoint, data for that timepoint was retrieved from (26). Finally, data on response to praziquantel treatment (Figure 2E) were retrieved from ERP002113. Single-cell RNA-Seq data from reference (32) was used to infer cell / tissue types impacted by drug treatment. Gene-set expression scores were calculated for gene sets (e.g., differentially expressed saposins, cathepsins, bax inhibitor-like proteins, and trematode eggshell proteins) using this dataset (GEO accession GSE146736). The Seurat object was imported into R, and the normalized RNA expression matrix and UMAP embeddings were extracted. For each gene set, normalized expression values were obtained by calculating gene-wise z-scores across all individual cells (subtracting the mean expression of each gene across cells and dividing by its standard deviation). A gene-set expression score was then calculated for each cell as the average of the z-scored expression values for all genes in the gene set and visualized on the published UMAP embedding. Analysis of glutamine synthase sequences was performed similar to the plexin domain-containing proteins noted above, aligning sequences with Clustal Omega and generating phylogenetic trees with IQ-TREE.

### InterPro domain enrichment

*S. mansoni* protein domain annotations were retrieved through BioMart on WBP19. To assess protein domains in gene products changing following treatment with both classes of drugs (gene products in DEG list of at least one TRP channel agonist and at least one antimalarial), enrichment testing was performed against the 8,058 genes tested in all four DESeq2 contrasts; testing was restricted to the 6,966 of these that carry at least one InterPro domain, so that genes without annotation could not contribute to either the foreground or the background.

Each InterPro domain represented by at least five genes in the universe and at least two genes in the tested set was evaluated with a one-sided Fisher exact test (alternative hypothesis: over-representation), and P values were adjusted within each direction by the Benjamini–Hochberg procedure. The micro-exon and plexin-containing families carry no InterPro domain, but were tested separately using the curated gene sets against the full 8,058-gene background. Fold enrichment is the proportion of the tested set carrying the domain divided by the proportion of the universe carrying it. Analyses used R 4.5.3 with tidyverse 2.0.0.

### Ca^2+^ flux assays

CHO-K1 cells were cultured in Dulbecco’s modified eagle medium supplemented with high glucose, GlutaMAX and pyruvate (ThermoFisher 10569010), MEM non-essential amino acids (ThermoFisher 11140050), 20mM HEPES, 10% fetal bovine serum (Gibco 16000-044) and penicillin (100 Units/mL) -streptomycin (100 µg/mL) at 37℃ and 5% CO_2_. *S. mansoni* TRP sequences (Smp_246790 and Smp_333650) were codon optimized (*Cricetulus griseus*, GenScript) and subcloned into pcDNA3.1(+). Plasmids were transiently transfected using Lipofectamine 3000 in T25 flasks, and replated onto poly-D-lysine coated black, clear bottom 96-well plates. Assays were performed the next day, replacing cell culture media with Fluo-4 Direct Calcium indicator in HBSS (ThermoFisher F10471), loading for 30 minutes at 37℃ followed by 30 minutes at room temperature. Assays measuring the activity of PZQ, MCLZ, MEF or ART were performed on a FlexStation3 (Molecular Devices). The same assay procedure was performed on untransfected cells control for activity at endogenous CHO-K1 pathways.

### Use of generative artificial intelligence disclosure

Claude (Anthropic) and ChatGPT (OpenAI) were used to assist with the development and modification of R scripts for data analysis and visualization, as well as manuscript proofreading and literature searches. The code used for these analyses is publicly available (https://github.com/zamanianlab/schistosome-antimalarial-rnaseq), enabling reproduction of computational pipelines. The authors retain full responsibility for the data analyses and manuscript content.

### Data availability statement

RNA-Seq data generated in this study are available in the NCBI Sequence Read Archive under BioProject PRJNA1544251. Re-analyzed datasets are available under BioProjects PRJNA597909, PRJNA1150591, PRJNA602528, PRJNA1177852 and PRJEB1281; life cycle expression data under ENA BioProjects PRJEB32839 and PRJEB30644; and single-cell data were retrieved from GEO accession GSE146736. The *S. mansoni* v10 genome and annotation (BioProject PRJEA36577) were obtained from WormBase ParaSite release 19.

## Acknowledgments

This work was supported by funding from NIH-NIAID R21 AI153545 (J.D.C and M.Z.). J.D.C. was supported by funding from the University of Wisconsin-Madison Global Health Institute. Schistosome-infected mice were provided by the NIH-NIAID Schistosomiasis Resource Center for distribution through BEI Resources, NIH-NIAID Contract HHSN272201700014I. The authors thank the UW-Eau Claire students in the fall 2025 BIOL 343 course, who contributed to initial QC and collaborative analysis of the RNA-seq data.

## List of abbreviations

ART: artemether
DEG: differentially expressed gene
DMEM: Dulbecco’s modified Eagle medium
DMSO: dimethyl sulfoxide
DyP: dye-decolorizing peroxidase
GOGAT: glutamate synthase
MCLZ: meclonazepam
MEF: mefloquine
MEG: micro-exon gene
PDG: plexin domain containing gene
PSI: plexin-semaphorin-integrin
PZQ: praziquantel
SmLE-PZQ-ER: Schistosoma mansoni LE PZQ-enriched resistant population
SmLE-PZQ-ES: Schistosoma mansoni LE PZQ-enriched sensitive population
TPM: transcripts per million
TRP: transient receptor potential
TRPM: transient receptor potential melastatin
UMAP: uniform manifold approximation and projection
WBP19: WormBase ParaSite release 19.

---

**Supplemental Figure 1.**
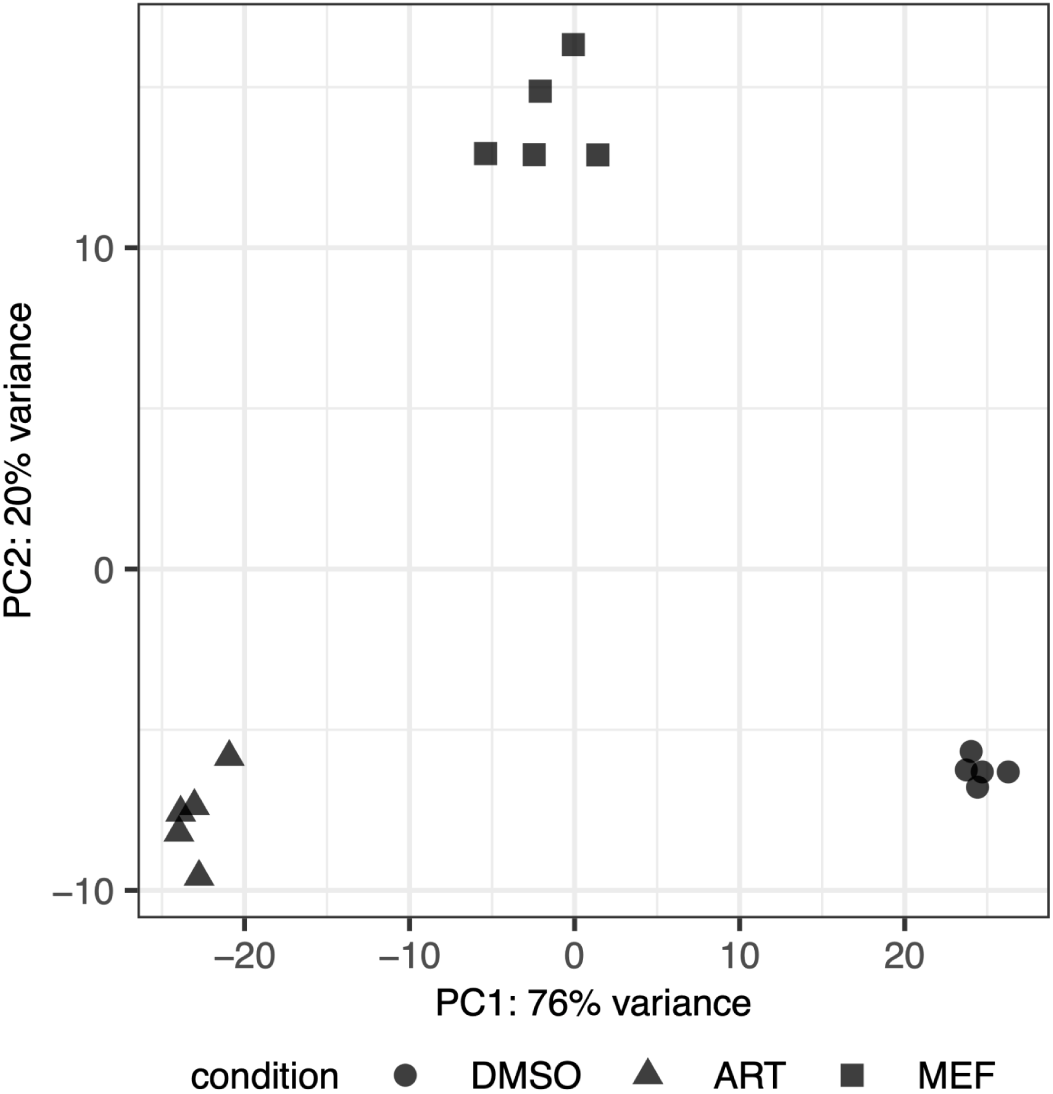
Principal component analysis of mefloquine and artemether treated cohorts. RNA-Seq data for DMSO control (circles), artemether (ART, triangles) and mefloquine (MEF, squares) treated cohorts.

**Supplemental Figure 2.**
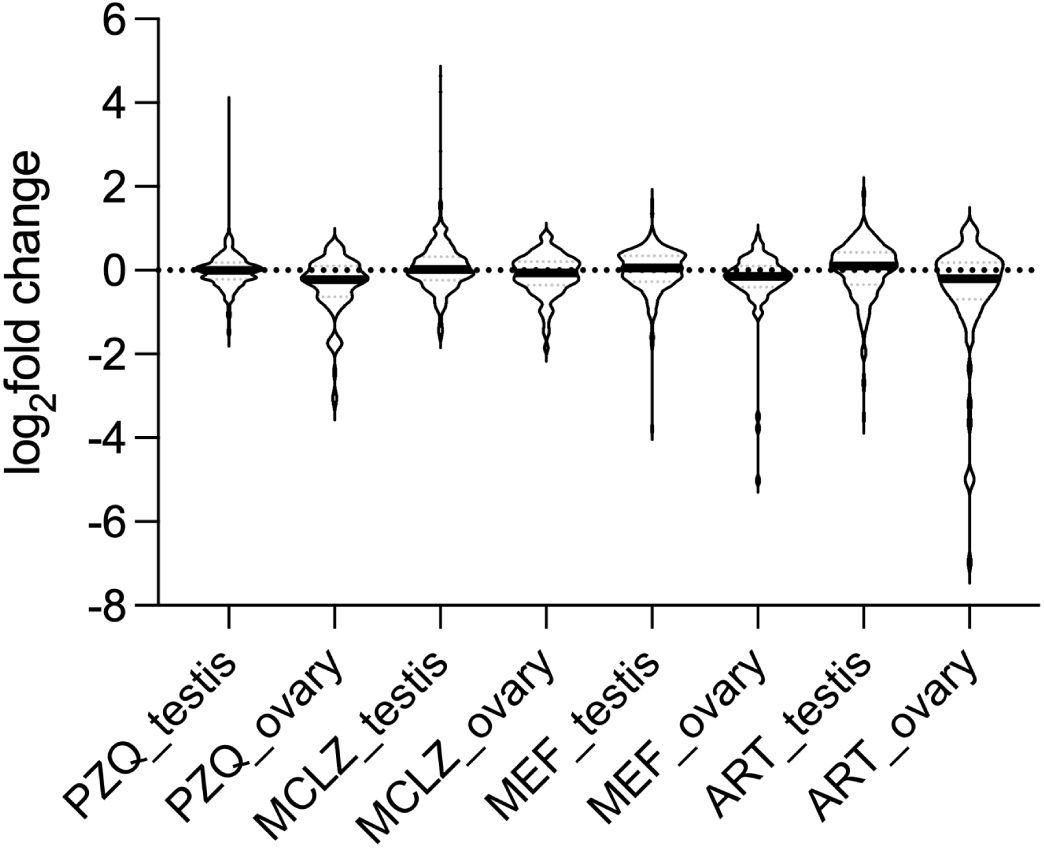
Expression of testis and ovary markers across drug treated samples. Expression of single-cell atlas markers (reference (32)) of late male germ cells (testis, n = 429 - 439) and late female germ cells (ovary, n = 84 - 85) relative to controls for each drug treatment. Bars reflect median log_2_ fold change values for each treatment.

**Supplemental Figure 3.**
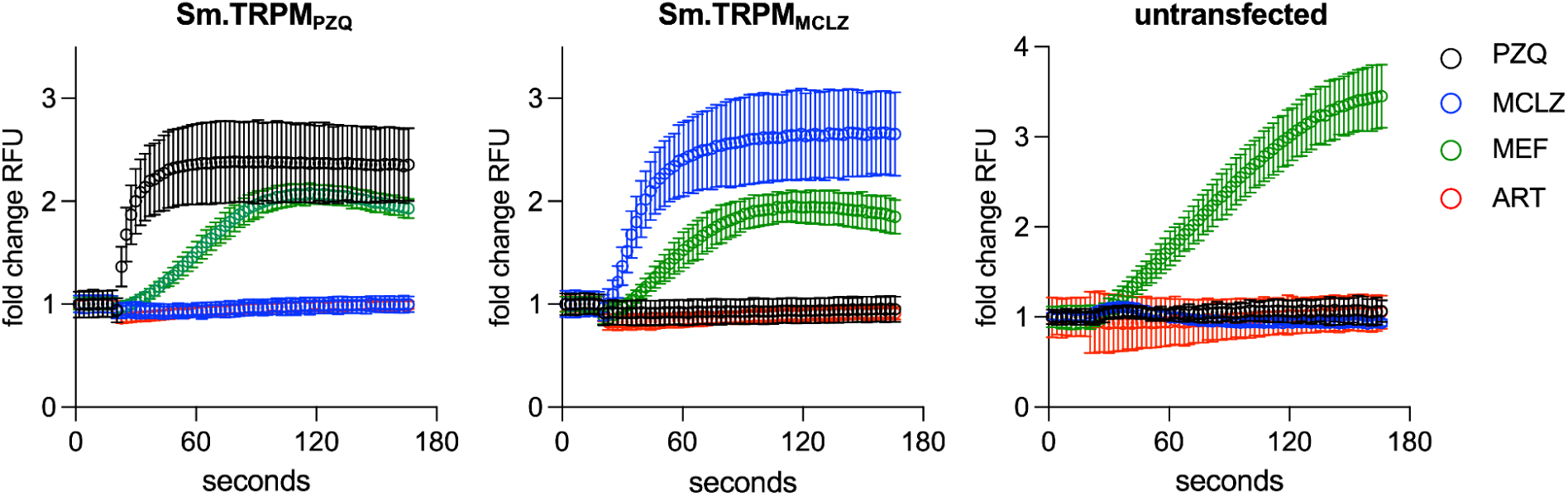
Activity of antimalarials artemether and mefloquine at schistosome TRP channels. Praziquantel (PZQ 10 µM, black), meclonazepam (MCLZ 10 µM, blue), mefloquine (MEF 30 µM, green) and artemether (30 µM, red) were applied to CHO-K1 cells transiently transfected with the target of PZQ (TRPM_PZQ_, left), the target of meclonazepam (TRPM_MCLZ_, middle), or untransfected cells (right). Cells were loaded with Flou-4 Direct Ca^2+^ Indicator, baseline readings were acquired for 20 seconds before compound addition, and Ca^2+^ flux is displayed by fold change in fluorescence (pre-compound addition / post-compound addition). ART was inactive in all conditions. MEF produced a Ca^2+^ signal in all conditions, including untransfected cells, indicating that this response is not due to actions on TRP channels.

**Supplemental Figure 4.**
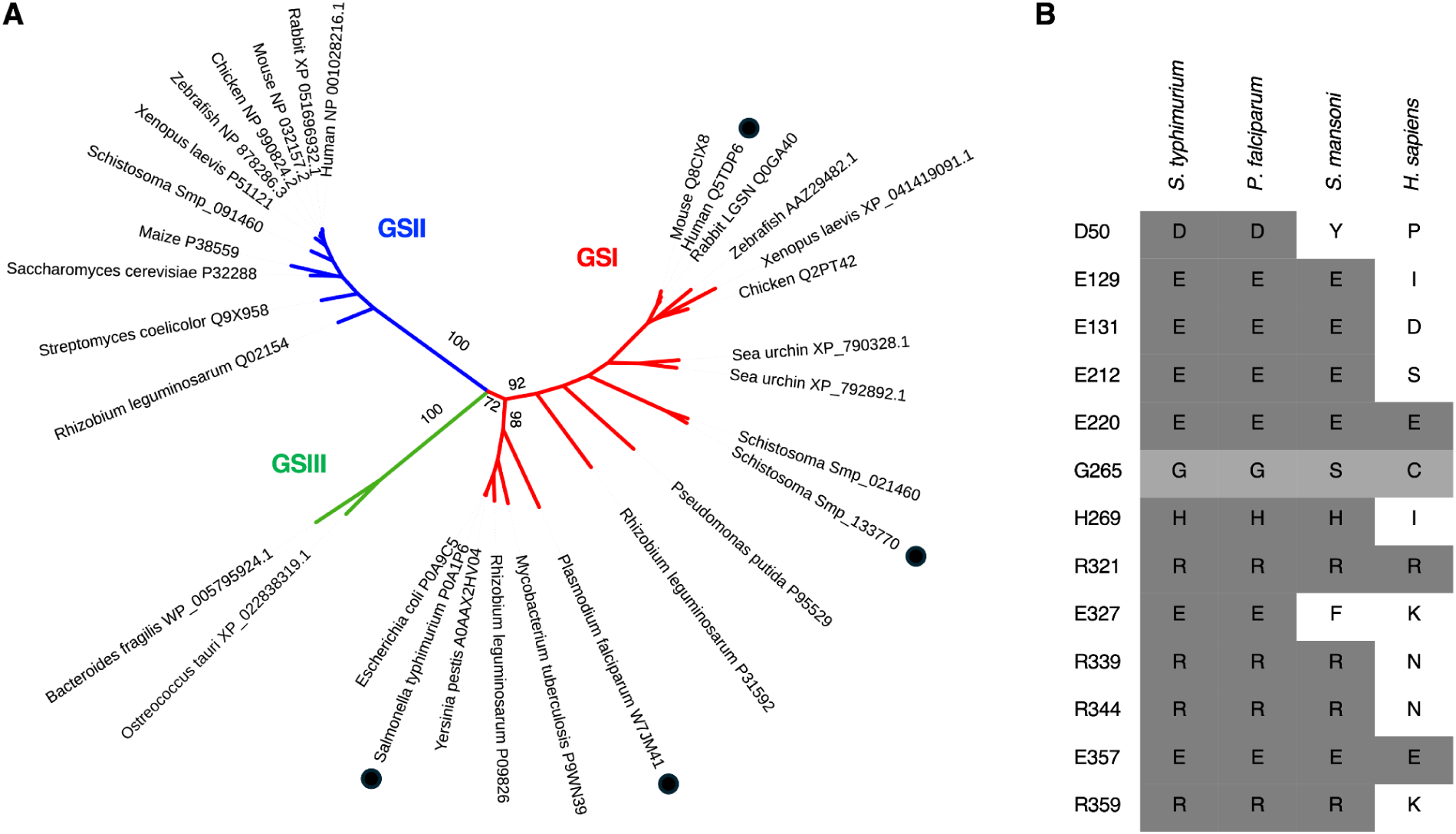
Analysis of glutamine synthetase-like sequences. **(A)** Unrooted maximum-likelihood tree of glutamine synthetase sequences. Bootstrap values are shown at key nodes. Red = class I glutamine synthetase sequences (GSI). Blue = class II glutamine synthetase sequences (GSII). Green = class III glutamine synthetase sequences (GSIII). Solid circles indicate species shown in greater detail in B. **(B)** Conservation of amino acids critical for GSI activity, based on structures in reference (92). Numbering reflects *Salmonella typhimurium* (P0A1P6) amino acid positions. Identical amino acids highlighted in dark grey. Light grey (G265) notes a position where the backbone carbonyl, and not the side chain, interacts with the glutamate substrate.

**Supplemental Table 1.** Change in cell and tissue specific markers with drug treatment. Percentage of markers from each cell type (identified in reference (32)) that are found in differentially expressed gene sets of various drug treatments; praziquantel (PZQ), meclonazepam (MCLZ), mefloquine (MEF) and artemether (ART).

|  | Markers (n) | PZQ | MCLZ | MEF | ART |
|---|---|---|---|---|---|
| early tsp−2+ | 134 | 7 | 16 | 7 | 17 |
| early vitellocytes | 153 | 21 | 17 | 31 | 39 |
| egc+ | 188 | 5 | 7 | 3 | 9 |
| eled+ neoblasts | 99 | 3 | 7 | 4 | 14 |
| esophageal gland | 26 | 0 | 69 | 81 | 77 |
| flame cells | 585 | 4 | 8 | 5 | 11 |
| GSC progeny | 231 | 14 | 14 | 10 | 23 |
| GSCs | 75 | 7 | 4 | 8 | 17 |
| gut | 81 | 5 | 11 | 15 | 86 |
| hes2+ | 73 | 5 | 10 | 1 | 10 |
| late female germ cells | 88 | 11 | 6 | 6 | 17 |
| late male germ cells | 449 | 4 | 7 | 5 | 10 |
| late vitellocytes | 243 | 28 | 18 | 42 | 49 |
| mature vitellocytes | 129 | 18 | 13 | 33 | 38 |
| meg−1+ | 408 | 3 | 7 | 2 | 6 |
| Mehlis' gland | 146 | 12 | 12 | 9 | 16 |
| muscle 1 | 310 | 12 | 5 | 5 | 14 |
| muscle 2 | 350 | 9 | 3 | 5 | 14 |
| muscle 3 | 238 | 10 | 3 | 7 | 14 |
| muscle 4 | 365 | 8 | 5 | 4 | 13 |
| muscle 5 | 330 | 8 | 5 | 6 | 14 |
| muscle 6 | 385 | 8 | 5 | 6 | 12 |
| muscle 7 | 352 | 9 | 3 | 6 | 13 |
| muscle 8 | 369 | 9 | 4 | 5 | 14 |
| neoblast 1 | 108 | 5 | 6 | 6 | 12 |
| neoblast 2 | 66 | 6 | 11 | 8 | 15 |
| neoblast progeny | 52 | 2 | 8 | 0 | 12 |
| neuron 1 | 278 | 6 | 5 | 2 | 8 |
| neuron 10 | 277 | 3 | 6 | 3 | 7 |
| neuron 11 | 298 | 6 | 7 | 5 | 12 |
| neuron 12 | 262 | 7 | 9 | 3 | 9 |
| neuron 13 | 319 | 4 | 4 | 4 | 11 |
| neuron 14 | 211 | 6 | 7 | 5 | 16 |
| neuron 15 | 266 | 5 | 8 | 3 | 11 |
| neuron 16 | 360 | 3 | 5 | 3 | 6 |
| neuron 17 | 268 | 6 | 4 | 4 | 10 |
| neuron 18 | 339 | 6 | 4 | 5 | 10 |
| neuron 19 | 235 | 3 | 7 | 1 | 8 |
| neuron 2 | 250 | 5 | 5 | 2 | 6 |
| neuron 20 | 351 | 6 | 5 | 4 | 12 |
| neuron 21 | 403 | 5 | 6 | 4 | 11 |
| neuron 22 | 130 | 4 | 3 | 4 | 15 |
| neuron 23 | 355 | 5 | 5 | 5 | 10 |
| neuron 24 | 285 | 5 | 6 | 3 | 10 |
| neuron 25 | 310 | 6 | 5 | 4 | 12 |
| neuron 26 | 237 | 5 | 6 | 3 | 7 |
| neuron 27 | 279 | 6 | 5 | 4 | 11 |
| neuron 28 | 269 | 5 | 3 | 2 | 7 |
| neuron 29 | 144 | 6 | 6 | 3 | 10 |
| neuron 3 | 330 | 5 | 4 | 3 | 8 |
| neuron 30 | 290 | 5 | 5 | 4 | 8 |
| neuron 31 (7b2−) | 178 | 7 | 6 | 3 | 12 |
| neuron 4 | 361 | 5 | 6 | 3 | 7 |
| neuron 5 | 430 | 4 | 5 | 4 | 7 |
| neuron 6 | 324 | 5 | 7 | 3 | 8 |
| neuron 7 | 220 | 4 | 7 | 2 | 10 |
| neuron 8 | 229 | 6 | 7 | 3 | 10 |
| neuron 9 | 321 | 5 | 7 | 2 | 9 |
| parenchyma 1 | 382 | 9 | 9 | 5 | 15 |
| parenchyma 2 | 411 | 7 | 7 | 3 | 11 |
| prom2+ | 103 | 7 | 16 | 2 | 25 |
| S1 | 89 | 7 | 7 | 8 | 16 |
| S1 progeny | 126 | 10 | 9 | 12 | 18 |
| sm13+ | 109 | 10 | 28 | 7 | 18 |
| tegument 1 | 375 | 11 | 39 | 9 | 20 |
| tegument 2 | 189 | 10 | 18 | 7 | 28 |
| zfp−1−1+ | 396 | 4 | 10 | 4 | 7 |

% of cluster markers that are DEGs
0 25 50 75 100

**Supplemental Table 2. Statistics and worm burden reduction following treatments following drug treatments shown in Figure 4**. Worms per mouse are shown as raw counts (mean ± standard error) for the groups named in the comparison columns. P value (exact, unadjusted) reflects result of two-sided exact Wilcoxon-Mann-Whitney test (shown in Fig 4). P value (Holm-adjusted) reflects Holm correction across the five treatment groups.

**Supplemental File 1. Mapped read counts and normalized transcript abundance (TPM) following drug treatment.** Reads from adult parasites treated with drug 7 weeks post infection and mapped to version 10 of the *S. mansoni* genome. For previously published datasets (praziquantel and meclonazepam), sample names are listed alongside the NCBI BioSample ID. **Sheet 1.** Praziquantel treatment read counts. Original data from reference (25), NCBI BioProject ID PRJNA597909. Data are re-mapped to the updated genome assembly. **Sheet 2.** Praziquantel treatment normalized transcript abundance in transcripts per million (TPM). **Sheet 3.** Meclonazepam treatment read counts. Data from reference (26). **Sheet 4.** Meclonazepam normalized transcript abundance in TPM. Data from reference (26). **Sheet 5.** Mefloquine and artemether treatment read counts. **Sheet 6.** Mefloquine and artemether treatment normalized transcript abundance in TPM.

**Supplemental File 2. Analysis of differential gene expression between control and drug treated samples.** Mapped read counts from Supplemental File 1 analyzed by DESeq2. **Sheet 1.** Praziquantel treatment. **Sheet 2.** Meclonazepam treatment. Data from reference (26). **Sheet 3.** Mefloquine treatment. **Sheet 4.** Artemether treatment.

**Supplemental File 3. Gene name and protein domain annotations.** Gene name and description (**Sheet 1**) and protein domains (PFAM and InterPro IDs, **Sheet 2**) were retrieved from WormBase ParaSite version 19 for *S. mansoni* (PRJEA36577).

**Supplemental File 4. Platyhelminth plexin-domain containing proteins.** FASTA file of plexin-domain containing protein sequences used for generating the phylogenetic tree in Figure 2C.

